# Discovering epistatic feature interactions from neural network models of regulatory DNA sequences

**DOI:** 10.1101/302711

**Authors:** Peyton Greenside, Tyler Shimko, Polly Fordyce, Anshul Kundaje

## Abstract

**Motivation:** Transcription factors bind regulatory DNA sequences in a combinatorial manner to modulate gene expression. Deep neural networks (DNNs) can learn the cis-regulatory grammars encoded in regulatory DNA sequences associated with transcription factor binding and chromatin accessibility. Several feature attribution methods have been developed for estimating the predictive importance of individual features (nucleotides or motifs) in any input DNA sequence to its associated output prediction from a DNN model. However, these methods do not reveal higher-order feature interactions encoded by the models.

**Results:** We present a new method called Deep Feature Interaction Maps (DFIM) to efficiently estimate interactions between all pairs of features in any input DNA sequence. DFIM accurately identifies ground truth motif interactions embedded in simulated regulatory DNA sequences. DFIM identifies synergistic interactions between GATA1 and TAL1 motifs from *in vivo* TF binding models. DFIM reveals epistatic interactions involving nucleotides flanking the core motif of the Cbf1 TF in yeast from in *vitro* TF binding models. We also apply DFIM to regulatory sequence models of *in vivo* chromatin accessibility to reveal interactions between regulatory genetic variants and proximal motifs of target TFs as validated by TF binding quantitative trait loci. Our approach makes significant strides in improving the interpretability of deep learning models for genomics.

**Availability:** Code is available at: https://github.com/kundajelab/dfim.

## 1. Introduction

Genome-wide biochemical profiling experiments have revealed millions of putative regulatory elements in diverse cell states. These massive datasets have spurred the development of deep neural network (DNN) models to predict cell-type specific or context-specific molecular phenotypes such as TF binding, chromatin accessibility and gene expression from DNA sequence^1 2 3^. Beyond high prediction accuracy, the primary appeal of DNNs is that they are capable of learning predictive sequence features and modeling non-linear feature interactions directly from raw DNA sequence without any prior assumptions. Hence, interpreting these purported black box models could reveal novel insights into the combinatorial regulatory code.

Advances in feature attribution methods for DNNs have enabled the identification of predictive cis-regulatory patterns in DNA sequences used as input to the models. Feature attribution methods estimate the contribution (or importance) of features, such as individual nucleotides or contiguous subsequences (e.g. motifs), in an input DNA sequence to a model’s output prediction. A perturbation-based, forward-propagation approach known as in-silico mutagenesis (ISM) quantifies the importance of a nucleotide in an input DNA sequence as the maximal change in the output prediction from the DNN model when the observed nucleotide (e.g. a G) at that position is mutated to any of the alternative bases (e.g. A, C or T). ISM has been used to score the potential molecular impact of genetic variants in regulatory DNA sequences^1 2 3^. However, ISM is computationally inefficient since each perturbation at every position in an input sequence requires a separate forward propagation to the output through the network.

**Figure 1:**
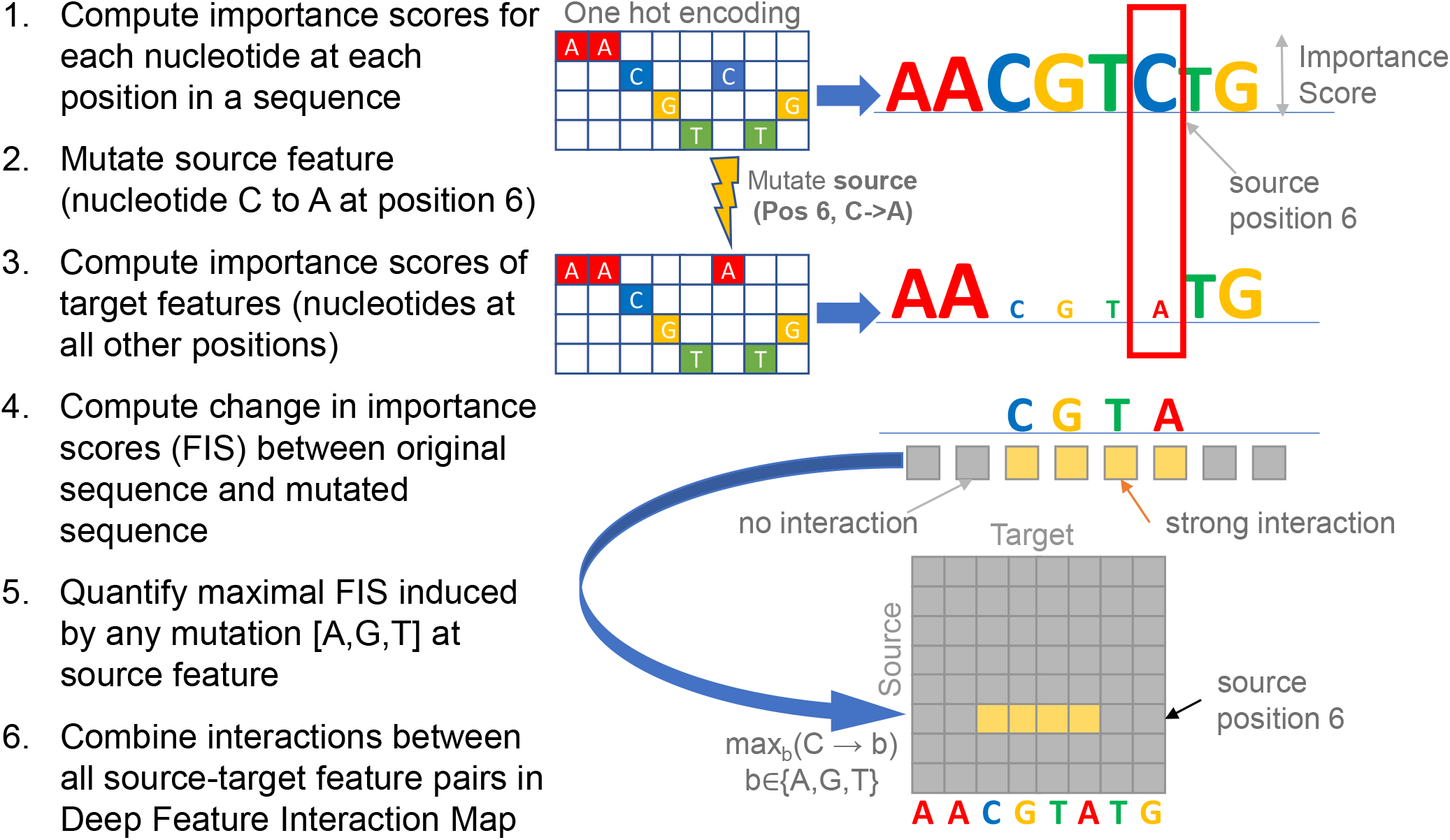
Deep Feature Interaction Maps: DFIM, illustrated in 6 steps, quantifies the maximal Feature Interaction Score (FIS) of every position in a sequence with all other positions.

ISM also fails to highlight important features masked by saturation due to buffering interactions with other features (e.g. multiple motif instances in a sequence that buffer each other)^4^. SHAP is a perturbation-based feature attribution method that borrows from game theory^5^. Max-Ent is a feature attribution method that uses a Markov chain Monte Carlo algorithm to find the maximum-entropy distribution of inputs that produced a similar hidden representation to the chosen input^6^. While SHAP and Max-Ent show improved sensitivity and specificity relative to ISM, they do not scale efficiently to comprehensively characterize feature importance across millions of regulatory sequences. An alternative family of computationally efficient back-propagation approaches decompose the output prediction corresponding to an input sequence by recursively propagating contribution scores through the layers of the DNN from the output to the input. One single backpropagation pass provides the contribution of all nucleotides in an input DNA sequence to the output prediction. The gradient of the output with respect to each nucleotide in the input DNA sequence - known as a saliency map7 - is one such estimate of importance and has been used to identify predictive nucleotides in regulatory DNA sequences. Other related approaches such as DeepLIFT^4^ and integrated gradients^8^ differ in the definition of the importance score that is backpropagated and provide improved sensitivity in the presence of saturation effects. DeepLIFT4 has also been shown to be an efficient approximation of SHAP scores^9^.

Current feature attribution methods only provide the importance of individual features. They do not highlight predictive, higher-order feature interactions that are learned by the DNN model. Perturbation-based approaches such as ISM cannot scale to comprehensively score all pairwise and higher-order interactions between nucleotides or subsequence features. Recently, an efficient algorithm was proposed to calculate SHAP-based pairwise feature interaction scores^9^ specifically from tree-based ensemble models. However, computing SHAP interactions from neural network models between all pairs of features in regulatory DNA sequences is computationally inefficient and cannot scale to reveal comprehensive interaction maps across millions of regulatory sequences.

Here, we present an efficient approach called Deep Feature Interaction Maps (DFIM) to estimate pairwise interactions between features (nucleotides or subsequences) in an input DNA sequence mapped to an associated regulatory phenotype by a neural network. We define a novel Feature Interaction Score (FIS) between any pair of features (source feature and target feature) in an input DNA sequence as the change in the importance score of the target feature when the source feature is perturbed, while keeping all the other features in the sequence intact. By leveraging efficient backpropagation-based feature attribution methods, we can efficiently compute FIS between all pairs of nucleotides or predictive motifs across large sets of input DNA sequence. Aggregate summary statistics of the pairwise Feature Interaction Score across multiple sequences provide insights into common, shared patterns of feature interactions.

We benchmark DFIM in controlled simulations that explicitly encode motif interactions. We use DFIM to reveal synergistic interactions between GATA1 and TAL1 motifs from *in vivo* TF binding models. We apply DFIM to reveal epistatic interactions involving nucleotides flanking the core motif of the Cbf1 TF in yeast from in *vitro* TF binding models. We also apply DFIM to regulatory sequence models of *in vivo* chromatin accessibility to reveal interactions between regulatory genetic variants and proximal motifs of target TFs as validated by TF binding quantitative trait loci.

## 2. Methods

We assume that we have trained a deep neural network to accurately map one-hot encoded DNA sequences *X* of length *L* to a categorical (binary or multiclass classification) or continuous (regression) output *O*. Let *Y* refer to the scalar predicted output *O* from the neural network for regression tasks. For classification tasks, let *Y* refer to the scalar input to the final output sigmoid (i.e. logit) of the neural network.

### 2.1. Nucleotide-resolution Feature Interaction Score (FIS)

We are given a one-hot encoded input DNA sequence *X*_0_ ∈ {0, 1}^{4×L}^ i.e. a matrix of size [4, *L*] such that *X*_0_[*b,p*] = 1 for the observed nucleotide *b* ∈ {*A, C, G, T*} at position 1 ≤ *p* ≤ *L* (Fig.1).

First, we compute *C*_*X*_0__ a matrix of size [4, *L*] that contains the importance (or contribution) of every nucleotide (rows) at each position in the sequence (Fig 1 Step 1). While our approach extends to any other efficient feature attribution method, for the analyses in this paper, we show results using both DeepLIFT4 and gradient saliency maps as importance scores^7^. In gradient-based saliency maps, for a specific input sequence *X*_0_, the output *Y*_0_ can be approximated by a first-order Taylor expansion *Y*_0_ ≈_*p,b*_ *w*_0_[*b, p*]*X*_0_[*b, p*] where *w*_0_ is the partial derivative (gradient) of *Y* with respect to the input sequence variable evaluated at *X*_0_ i.e. 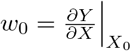. It is worth noting that the entire gradient matrix *w*_0_ can be computed efficiently in one backpropagation pass. We then perform a point-wise multiplication of the gradient matrix *w*_0_ with the one-hot encoded observed input sequence *X*_0_ to obtain the importance scores for the observed nucleotides *b* at each position *p* i.e. *C*_*X*_0__ = *w*_0_[*b, p*]*X*_0_[*b, p*]. Only the observed nucleotides at each position can have non-zero values. DeepLIFT contribution scores quantify the sensitivity of the output to finite changes in the input^4^. This is in contrast to gradients, which measure the sensitivity of the output to infinitesimal changes in the input. Specifically, the DeepLIFT algorithm backpropagates a score (analogous to gradients) which is based on comparing the activations of all the neurons in the network for the actual input sequence *X*_0_ to those obtained when using neutral “reference” sequences^4^. We use dinucleotide-shuffled versions of *X*_0_ as reference sequences unless otherwise specified.

Our goal is to query the neural network to estimate the interaction between the observed nucleotide at one position in the sequence (source feature) and the observed nucleotide at some other position (target feature) in the sequence. Let (*α, s*) represent the source feature i.e. the observed source nucleotide *α* ∈ {*A, C, G, T*} at a source position s such that *X*_0_[*α, s*] = 1. Let (*β, s*) represent the target feature i.e. observed target nucleotide *β* ∈ {*A, C, G, T*} at some target position *t* such that *X*_0_[*β, t*] = 1.

Intuitively, we define the Feature Interaction Score *FIS*_*X*_0__((*β, t*)|(*α, γ, s*)) of the target feature on the source feature as the change in the importance score of the target feature (*β, s*) when the source feature (*α, s*) is mutated to a different nucleotide (*γ, s*). To compute FIS, we create a new mutated sequence 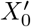 from *X*_0_ where we switch the observed nucleotide *α* at source position *s* to a different mutant nucleotide *γ* ∈ {*A, C, G, T*}, while keeping the nucleotides at all other positions as they were in *X*_0_ (Fig 1 Step 2). We then compute the importance matrix *C*_*X*_0__′ for 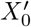 as we did for *X*_0_ (Fig 1 Step 3). The *FIS* of the target feature with the source feature is defined as

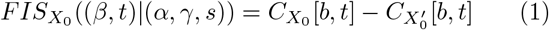

Since, only two backpropagation passes are required to compute *C*_*X*_0__ [, *t*] and 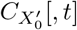 for all 1 ≤ *t* ≤ *L*, we can efficiently compute the *FIS* of all target features *FIS*_*X*_0__ (*|(*α, γ, s*)) in a sequence with respect to a specific source feature mutation (Fig 1 Step 4). Note that the *FIS* is a directional interaction score of the target with the source. In some cases, we may only be interested in the magnitude of the score rather than its sign. In such cases, we use the absolute value of the *FIS*.

We define the maximal Feature Interaction Score (*maxFIS*) of the target feature with the source feature as the maximal *FIS* marginalized over all possible values of the mutant nucleotide *γ* at the source feature (*α, s*) i.e *maxFIS*_*X*_0__((*β,t*)|(*α, s*)) = *max_γ_*((*β,t*)|(*α,γ, s*)) (Fig 1 Step 5).

A nucleotide-resolution Deep Feature Interaction Map (DFIM) summarizes the *maxFIS* scores for all pairs of source and target features in an input DNA sequence (Fig 1 Step 6).

### 2.2. Aggregate statistics of nucleotide-resolution FIS over multiple input sequences

In order to analyze the prevalence of the *FIS* between a source position *s* and target position *t* across a collection of input sequences *X_i_*, we first identify the subset of sequences *S* = {*X_i_*} that have identical source nucleotides at the source position and identical target nucleotides at the target position i.e ∀*X_i_*, *X_j_* ∈ *S*, *X_i_* [*α, s*] = *X_j_* [*α, s*] = 1 AND ∀*X_i_, X_j_* ∈ *S,X_i_*[*β, s*] = *X_j_*[*β, s*] = 1. We then compute aggregate statistics such as the mean of the *FIS* or absolute *FIS* corresponding to each ((*β, t*) |(*α, γ, s*)) over all sequences in the subset *S*. (See Fig 8 as an example).

### 2.3 Motif-resolution Feature Interaction Score

We are often interested in the *FIS* of a specific target motif {(*β_p_, t_p_*),…, (*β_q_, t_q_*)} i.e. a specific subsequence of nucleotides {*β_p_*,…,*β_q_*} at a specific subset of contiguous target positions {*t_p_*,…,*t_q_*} with a source nucleotide-resolution feature (*α, s*) (i.e. specific source nucleotide at specific source position) such as a regulatory single nucleotide variant (SNV). In such a case, we compute the *FIS* of a target motif with a source nucleotide feature as the difference of the sum of importance scores across all target nucleotides {(*β_p_, t_p_*),…, (*β_q_, t_q_*)} in the target motif in the original sequence *X*_0_ and the mutated sequence 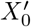 (obtained by mutating (*α, s*) in *X*_0_ to (*γ, s*)).

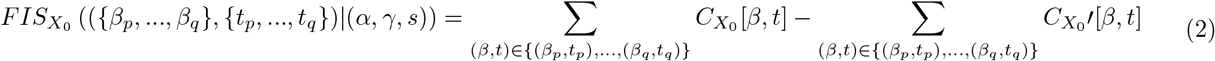

To compute the *FIS* of a target motif {(*β_p_,t_p_*),…, (*β_q_,t*_q_)} with a source motif {(*α_k_,t_k_*),…, (*α_l_,t_l_*)} (See Fig 3 as an example), we use a different source mutation method. One option would be use the maximal *FIS* of the target motif over all possible single nucleotide mutations of each position in the source motif. However, this procedure is computationally infeasible for long motifs. We instead, generate one mutant sequence, where we mutate the one-hot encoding (where rows 1-4 correspond to A,C,G,T) of all positions {*s_k_*,…, *s_l_*} in the source motif to the expected background GC nucleotide frequency *f_GC_* i.e. the mutant sequence 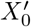 has 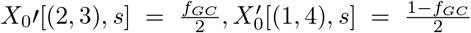. The *FIS* of the target motif with the source motif is once again the difference of the sum of importance scores across all target nucleotides {(*β_p_,t_p_*),…, (*β_q_,t_q_*)} in the target motif feature between the original sequence *X*_0_ and the mutated sequence 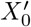.

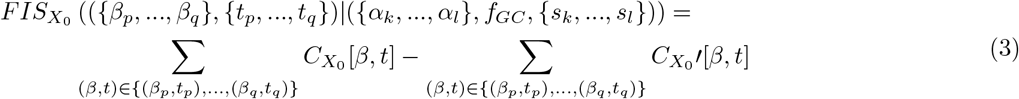

### 2.4. Statistical significance of FIS

Given a continuous distribution of *FIS*, across a collection of input sequences, we define statistically significant interactions based on an empirical null distribution of scores from dinucleotide shuffled versions of the input sequences. For each dinucleotide shuffled input sequence, we compute FIS for all nucleotide pairs. We fit a Gaussian distribution to this null empirical distribution of *FIS* scores. *FIS* values passing a *p*-value of 0.05 with respect to this null distribution are considered statistically significant. We use the Benjamini-Hochberg procedure for multiple hypothesis correction. SFig 1 demonstrates how the null model can be used to identify responding motifs in the context of a longer sequence.

### 2.5. Comparison of DFIM to SHAP Interaction Scores and pairwise ISM interaction scores

For an input sequence with *F* features (nucleotides/motifs), SHAP interaction scores scale at least quadratically to compute all pairwise interactions giving a complexity ranging from *O*(*F*^2^) to *O*(2^*F*^)^9^. A pairwise ISM-based interaction score, defined as the difference between the ISM score obtained by jointly mutating two features and the sum of the ISM scores of individual features, also has a complexity of *O*(*F*^2^). For DFIM, we require one backpropagation pass to obtain importance scores for the original sequence. Then for each of the *F* source features, we need one more backpropagation pass to obtain *FIS* of that source with all target features. Thus, DFIM exhibits a complexity of *O*(*F*) scaling linearly in the number of features. Our proposed *FIS* is essentially an efficient approximation of SHAP interaction scores. Further, in contrast to SHAP interaction scores and pairwise ISM interaction scores which are necessarily symmetric over the source and target, *FIS* is directional and can produce asymmetric interaction scores.

**Figure 2:**
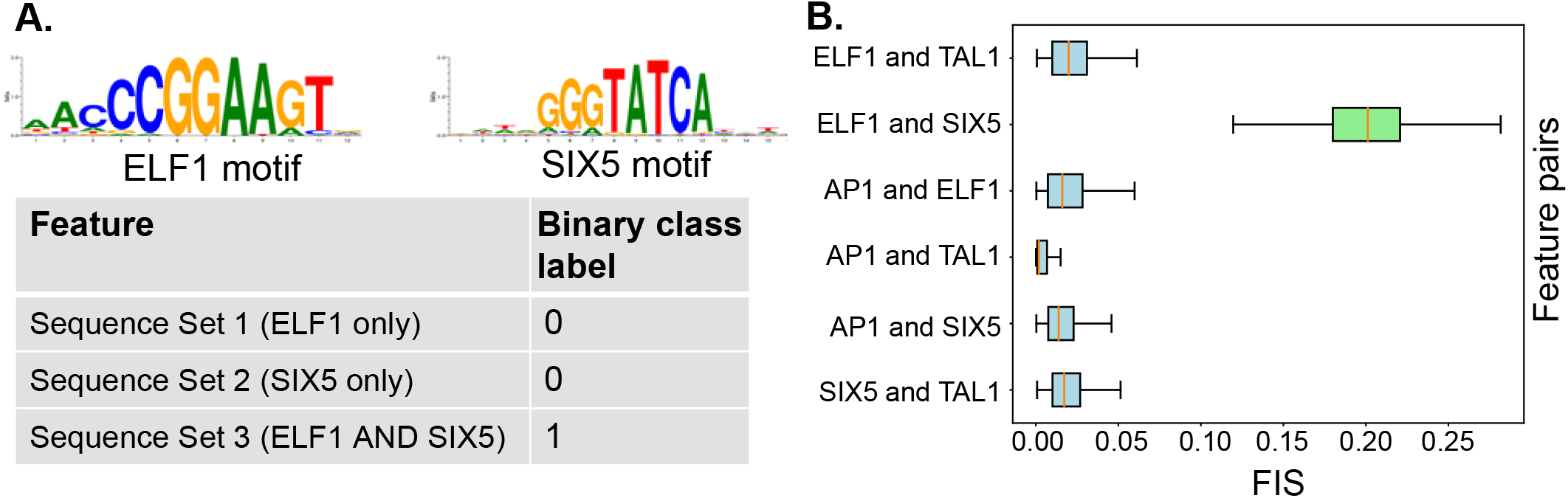
**A.** Simulated dataset: Sequences in the positive class contain both ELF1 and SIX5 motif instances. **B.** Distribution of feature interaction scores (FIS) for different motif pairs. Pairs of ELF1 and SIX5 motifs are the only pair with high FIS.

## 3. Results

### 3.1. Benchmarking FIS on ground-truth motif interactions embedded in simulated regulatory DNA sequences

To benchmark *FIS*, we simulated 60K random DNA sequences (0.46 G/C frequency) of length 200 bp. We divided these into 3 sets of 20K sequences. We randomly embedded 1 or 2 instances of the ELF1 motif (using the highest affinity sequence from Position Weight Matrix^10^) in the sequences in Set 1, 1 or 2 instances of the SIX5 motif in Set 2 and 1 or 2 instances of both ELF1 and SIX5 motifs in Set 3. We further independently embedded 0 or 1 instances of the AP1 and TAL1 motifs in a random subset of sequences across all 3 sets10 (Supp. Methods). We then set up a binary classification task where all sequences in Set 3 (ELF1 and SIX5) were labeled as positive and all other sequences from Sets 1 and 2 were labeled as negatives (Fig 2A). We trained a Convolutional Neural Network (CNN) with one convolutional and one dense layer (Supp. Methods). We achieved 100% classification accuracy on held out validation set of sequences indicating the model had learned the necessary interaction between ELF1 and SIX5. We computed motif-resolution *FIS* for all pairs of embedded motif instances (SIX5, ELF1, AP1 and TAL1) for all sequences in the positive class (i.e. Set 3). We used DeepLIFT with a fixed GC reference for computing importance scores since the underlying sequences were generated using a fixed GC background. Only pairs of SIX5 and ELF1 motifs (positive control) showed strong *FIS* (Fig 2B, green distribution), compared to all other pairs of motifs (negative controls) demonstrating that can effectively discriminate ground truth interactions learned by a neural network. We further assessed the significance of these interactions using a empirical null distribution from dinucleotide shuffled sequences and found that the vast majority of true ELF1-SIX5 interactions have significant (*p* < 0.05) *p*-values, even after multiple hypothesis correction. None of the other motif pairs show statistically significant interactions (SFig 2A,B). The results are replicated using gradient saliency maps as importance scores (SFig 2C,D).

### 3.2. Uncovering epistatic motif interactions of co-binding TFs from CNN models of *in vivo* TF binding

We analyzed CNN models of *in vivo* TF binding to investigate epistatic interactions between motifs of co-binding TFs. We trained a multi-task CNN model to classify 1 kbp sequences centered at GATA1, GATA2 and TAL1 ENCODE ChIP-seq peaks (positive class) in erythroid K562 cells from all other chromatin accessible DNase-seq peaks in K562 (negative class)^11^>12 (Supp. Methods). The CNN model with 5 convolutional layers (25 convolutions, size 10), a max pooling layer (size 25) and a sigmoid activation (Supp. Methods), achieved mean auROC of 0.953 and mean auPRC of 0.459 across all three tasks on held-out test set. Next, we identified all matches to the known motifs of GATA1 and TAL1 in all ChIP-seq peak sequences (Supp. Methods). We then computed motif-resolution *FIS* (using DeepLIFT with shuffled reference as importance scores) for all pairs of GATA1, TAL1 motif instances across all sequences using GATA1 as the source motif. We observed several instances with strong *FIS* between proximal GATA1 and TAL1 motifs which corroborates their experimentally validated co-binding interactions13 (Fig. 3A). To understand the relationship between the distance between motif instances and their interaction scores, we binned GATA1 and TAL1 motif pairs into 4 distance bins - within 20bp (n=13,004), 20-50bp (n=18,898), 50-100bp (n=28,684), and 100-200bp (n=21,1154). We compared the distribution of *FIS* for the motif pairs across the bins. As expected, TAL1 and GATA1 motifs in close proximity (<20 bp) showed statistically significant higher interaction scores than all three other bins (*p*<1e-16, Mann Whitney test for all 3 comparisons) (Fig. 3A). However, interestingly, we observed some strong long-range interactions between motifs as far as 70 bp apart (Fig. 3B), an observation corroborated by a recent analysis of SNP effects on TAL1 ChIP-seq signal in erythroid cells that found that GATA1 motif mutations impact TAL1 binding at distances as great as 75 bp^14^. The interactions were also symmetric, such that mutating TAL1 demonstrated a similar distribution of *FIS* on GATA1 (SFig. 4).

**Figure 3:**
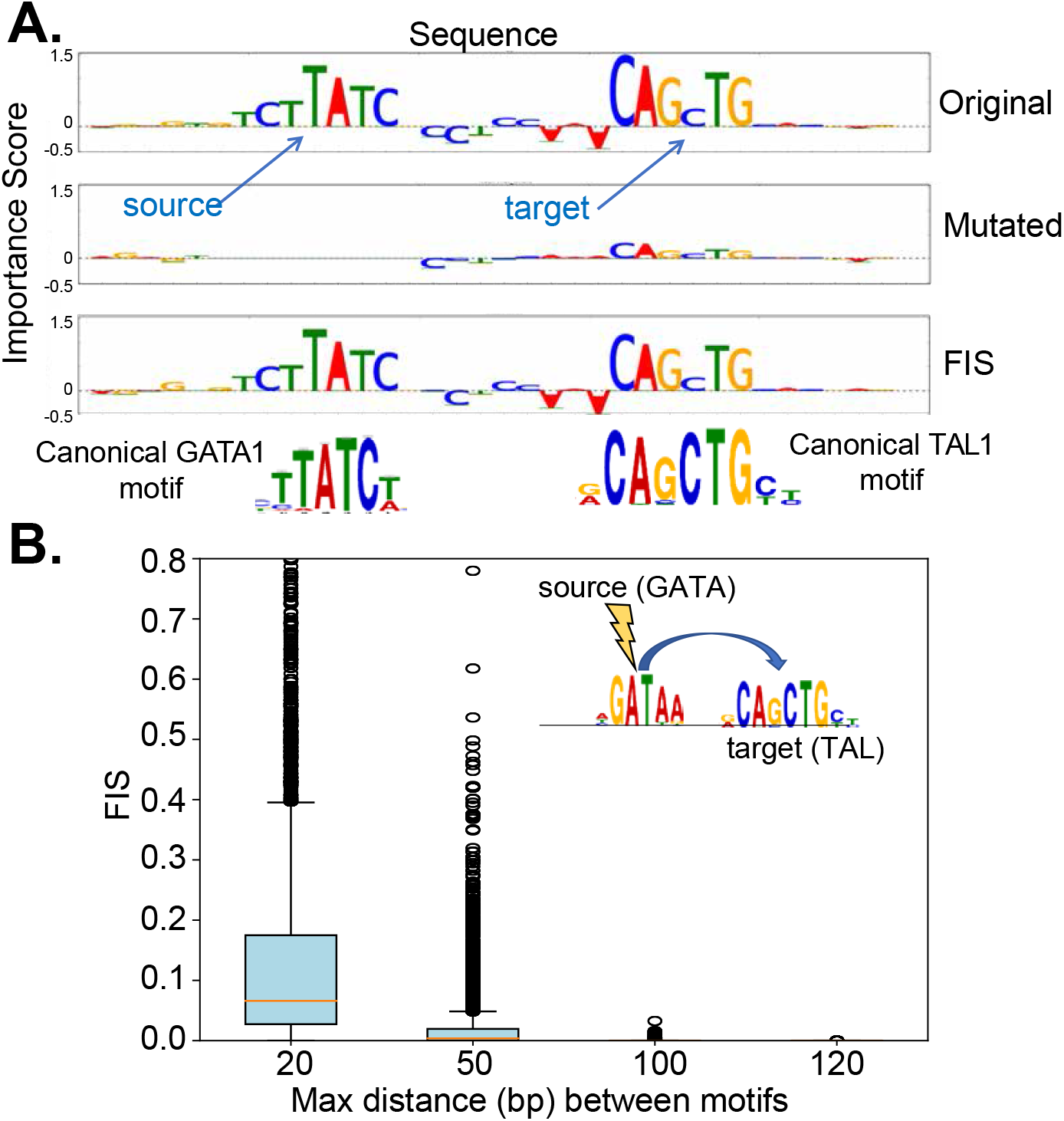
**A.** Example sequence showing interaction between GATA and TAL motif. Top row shows the importance scores for the original sequence. When the source GATA motif is mutated, the feature importance of the target TAL1 motif drops (second row) indicating a strong interaction between the two motifs (high *FIS*) (third row). **B.** Distribution of *FIS* scores for GATA1-TAL1 motif pairs stratified by relative distance between motifs. GATA1 motifs exhibit strong interactions with TAL1 motifs that are within 20bp distance.

### 3.3. Discovering interactions between regulatory variants and their target TF motifs from CNN models of *in vivo* chromatin accessibility

DNNs mapping regulatory DNA sequences to TF binding and chromatin accessibility have been previously used to score the predicted in-silico allelic effects of putative regulatory genetic variants based on ISM^1 2 3^. Here, we instead use *FIS* to investigate an orthogonal question - What proximal sequence features are affected by (interact with) regulatory genetic variants? Tehranchi *et al*. developed a pooling-based approach to identify thousands of SNVs that have allelic effects on TF binding (as measured by ChIP-seq) across a large collection of genotyped lymphoblastoid human cell-lines^15^. They provide coordinates, effect sizes, reference/alternative alleles and the allele with stronger binding for statistically significant binding QTLs (bQTLs) and non-significant background control SNVs in ChIP-seq peaks for JUND, NFKB, SPI1, STAT1 and POU2F1. This dataset provides an excellent resource to investigate the feature interactions of bQTLs. Further, we wondered if we could discover bQTL feature interactions for different TFs from a single DNN model trained to predict chromatin accessibility (instead of TF binding) from sequence.

**Figure 4:**
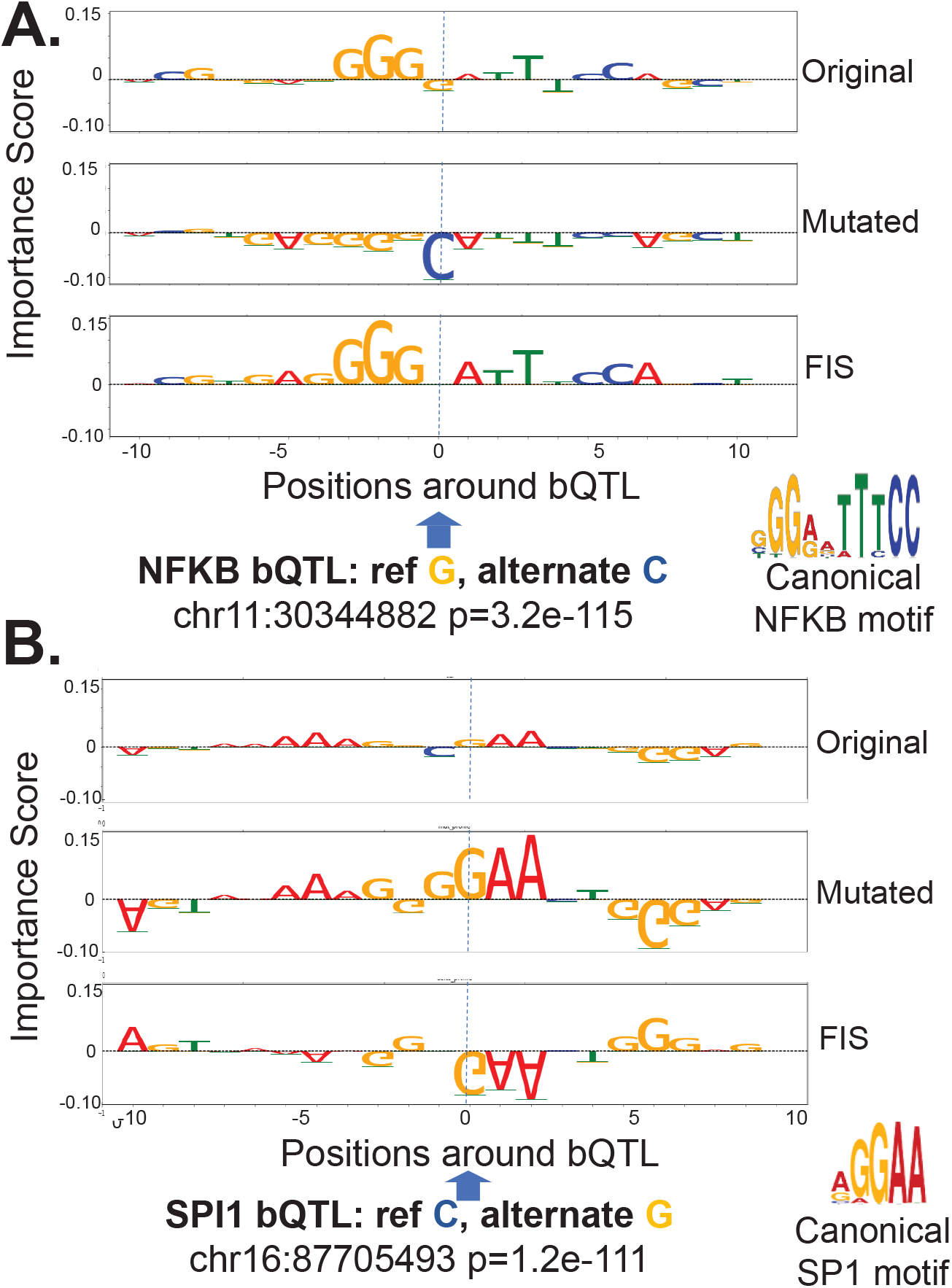
Examples of an NFKB bQTL (**A.**) and a SPI bQTL (**B.**) exhibiting strong interactions (FIS) with nucleotides in overlapping motif instances. Top and second row in both panels are the feature importance scores of each nucleotide around the bQTL for the reference and alternate allele respectively. The third row in both panels is the feature interaction score (*FIS*) indicating the change in importance when the reference allele is mutated to the alternate allele. For both bQTLs shown, the G allele is predicted to favor binding (positive importance scores of nucleotides in overlapping motif instance). The G allele is also the measured stronger binding allele.

Hence, we trained a multi-task (18 tasks) CNN model to map 1kbp length DNA sequences to binary chromatin accessibility profiles across 16 primary hematopoietic cell types (with ATAC-seq data)16 and 2 ENCODE cell-lines (with DNase-seq data) including the GM12878 lymphoblastoid cell-line (LCL)^12^. The model achieved high performance on the test set (average auPRC = 0.69, auROC=0.91). We used the LCL task to investigate bQTL feature interactions using (DeepLIFT with shuffled reference as importance score). We restricted our analysis to the statistically significant (allelic binding *p*<5e-05 as recommended by Tehranchi *et al*.^15^) bQTLs that overlapped the DNase-seq peaks in GM12878.

**Figure 5:**
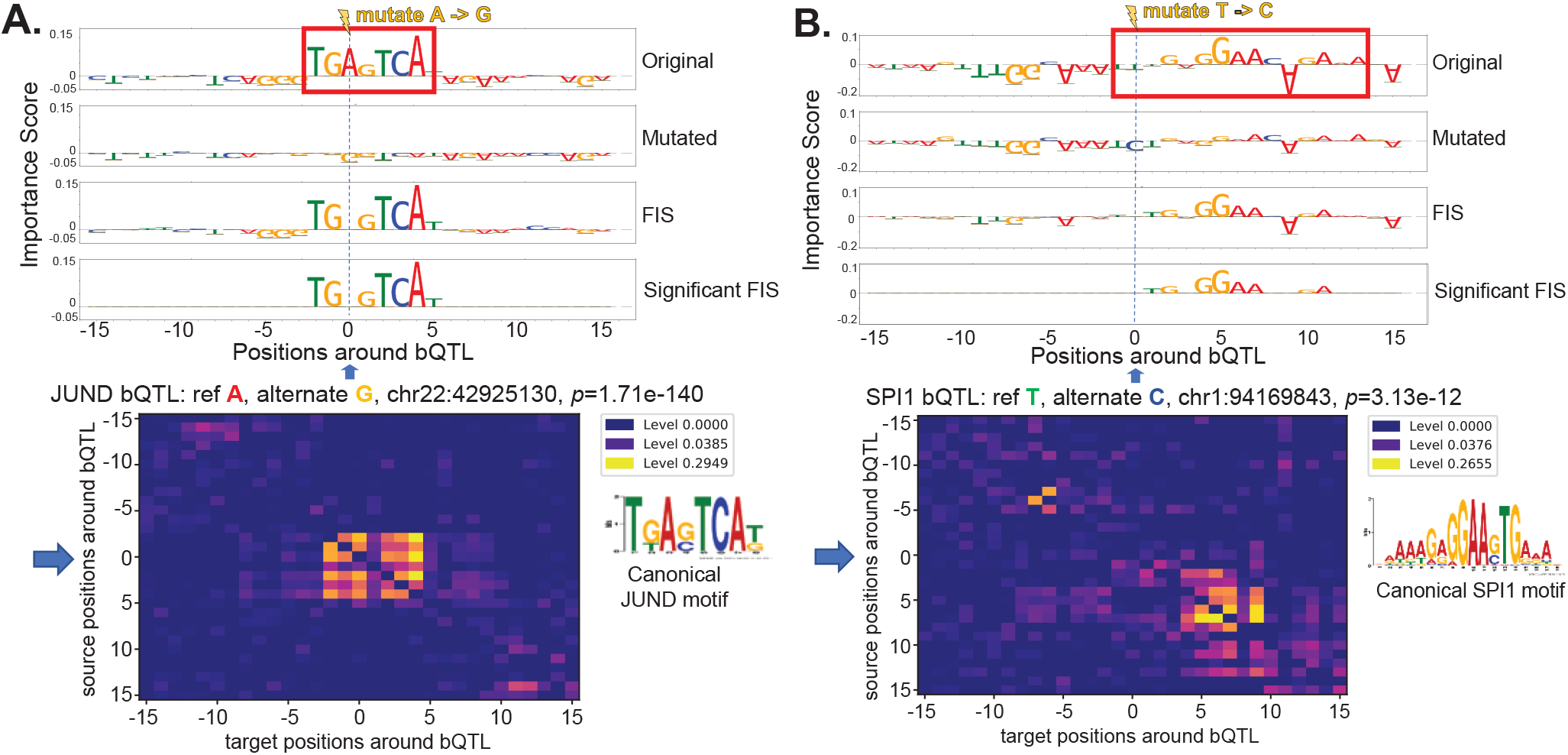
Examples of NFKB bQTL (**A.**) and JUND bQTL (**B.**) exhibiting strong feature interaction scores (*FIS*) with nucleotides in an overlapping motif and flanking motif respectively. Row 1 and 2 in both panels are the feature importance scores of each nucleotide around the bQTL for the reference and alternate allele respectively. Row 3 in both panels show the feature interaction scores (*FIS*) indicating the change in importance when the reference allele is mutated to the alternate allele. Row 4 shows only statistically significant interactions. Row 5 shows the deep feature interaction map as a heatmap.

To understand proximal interactions, for each bQTL, we used *FIS* to estimate the effect of mutating the reference allele to all alternate alleles at the source QTL on every target nucleotide +/- 15 bp around the QTL. First, we observed strong positive (Fig 4A) and negative (Fig 4B) interactions of bQTLs with nucleotides of overlapping target TF motifs. The direction of the allelic effect (stronger or weaker ChIP-seq signal) of the reference and alternate bQTL alleles on TF binding also matched the predicted direction of change (stronger or weaker motif score) E.g. A significant JUND bQTL at chr22:42925130 falls in a high affinity JUND binding motif (Fig 5A). The reference A allele has higher binding than the alternative G allele with *p*-value 1.71e-140 in the Tehranchi *et al.* study^15^. *FIS* predicts that the G allele (weaker allelic binding) but not the A allele (stronger allelic binding) will destroy the importance of the entire JUND motif.

Next, we also found several TF-bQTLs in the flanking nucleotides of weak affinity motif matches of the target TF having significant interaction effects with the entire motif. E.g. a significant SPI1 bQTL at chr1:94169843 has reference allele T (with stronger binding) and alternate allele C. The bQTL is in the flanking nucleotides of a low affinity SPI1 site where only the core GGAA matches the canonical motif. *FIS* predicts that the C allele (weaker binding) destroys the importance scores of the core GGAA element (Fig 5A). Tehranchi *et al*.^15^ and several other studies have reported that a large fraction (70-90%) of QTLs do not overlap high affinity instances of canonical TF motifs. We hypothesize that several QTLs may be affecting flanking nucleotides of weak affinity TF motif instances. Finally, while most bQTLs with statistically significant interactions exhibit the maximal absolute interaction with other nucleotides within 10 bp of the bQTL, we also observe strong and significant longer-range interactions at distances ranging from 20-200 bp (Fig 6A) e.g. an SPI1 bQTL has a significant interaction with a proximal SP1 motif but also a strong interaction with a RUNX1 motif 20 bp away (Fig 6B). SPI1 QTLs were also found to affect motifs 100s of base pairs away (SFig 5).

As a negative control, for each TF, we also evaluated the *FIS* of a matched number of conservative control SNVs from the Tehranchi *et al*.^15^ study that overlap the TF’s ChIP-seq peaks and LCL DNase-peaks with least significant allelic effects on binding (allelic binding *p* ≈ 1). For each bQTL and control SNV, we recorded its maximal abso lute *FIS* (*maxAbsFIS*) over all target nucleotides +/-15 bp around the SNV. For all the TFs, we found that the bQTLs exhibit significantly (Mann Whitney test) stronger *maxAbsFIS* than control SNVs (Fig 7), indicating that FIS may be an alternative approach to ISM to identify putative regulatory variants. This result was replicated using gradient based importance scores (SFig 3).

**Figure 6:**
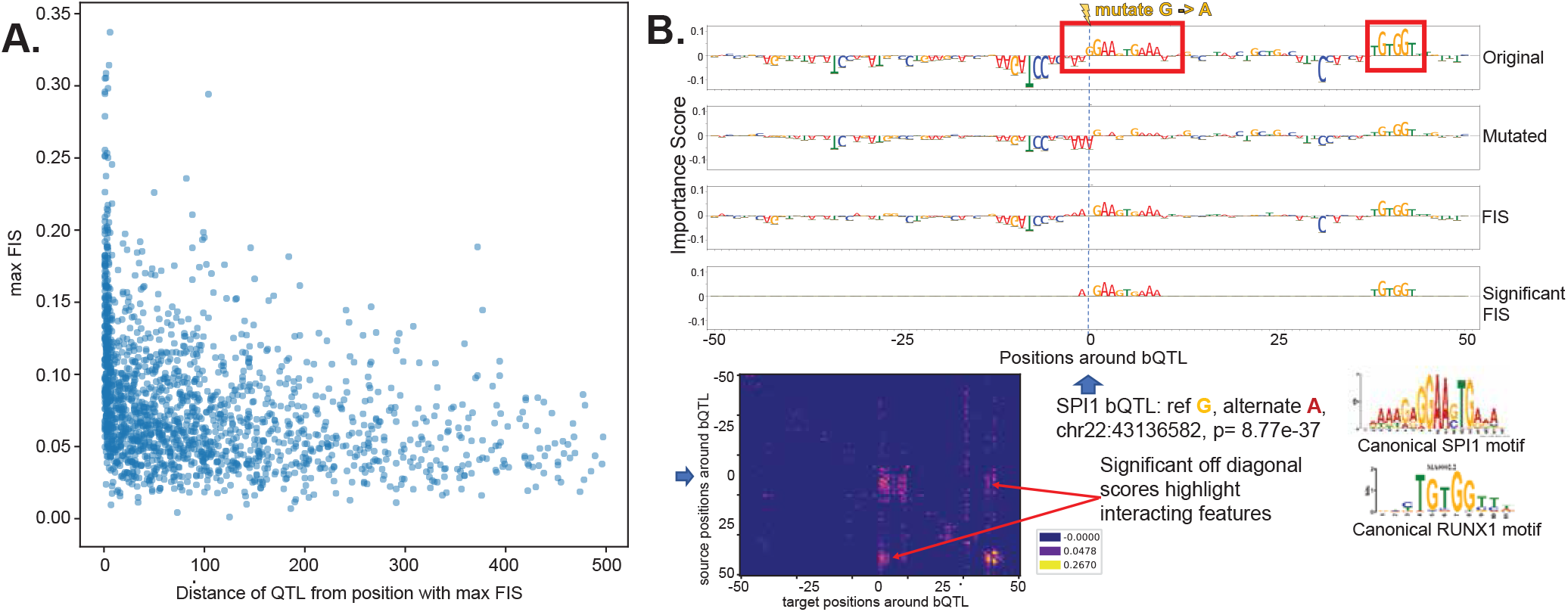
**A.** Maximum absolute feature interaction scores (*FIS*) (y-axis) as a function of distance between SPI1 bQTL and maximally interacting nucleotide within 1 kbp of the bQTL. **B.** Example of a SPI1 bQTL showing significant interactions with an overlapping SP1 motif and a proximal RUNX1 motif 40bp from the bQTL. Row 1 and 2 show the feature importance scores of each nucleotide around the bQTL for the reference and alternate allele respectively. Row 3 in both panels show the feature interaction scores (*FIS*) indicating the change in importance when the reference allele is mutated to the alternate allele. Row 4 shows only statistically significant interactions. Row 5 shows the deep feature interaction map as a heatmap.

### 3.4. Discovering interactions between nucleotides flanking the core sequence motif of the Cbf1 TF in yeast from *in vitro* binding DNN models

Paralogous TFs have been recently shown to have distinct sequence affinity preferences to nucleotides flanking the core canonical binding motifs. Le and Shimko *et al.* recently developed a microfluidics based *in vitro* TF binding assay called BET-seq to investigate this question^17^. They used the BET-seq assay to measure high-resolution *in vitro* binding affinity landscapes of the yeast TFs Cbf1 and Pho4 to a high complexity library of > 1 million DNA sequences with a fixed central core E-box sequence (CACGTG) and 5 variable flanking nucleotides on either side. They trained a feed forward neural network to predict relative binding affinity (Δ Δ G) for each of the TFs from the 10bp flanking sequences (using a flattened one-hot encoding) in the library17 (Fig 8 A). The model architecture consisted of 3 dense layers of sizes 500, 500 and 250 with ReLU activation followed by batch normalization and dropout (p=0.25) with a final dense classification layer having a linear activation. They used a distillation approach to interpret the NN model by fitting a linear model with all mononucleotide features across all positions and all dinucleotide features across all pairs of positions to the output predictions of the NN. They found that dinucleotide features were critical for the linear model to have a good fit (*r*^2^ > 0.95) especially for Cbf1. They then estimated the contributions of all pairwise interaction terms by comparing the mononucleotide+dinucleotide linear model to a mononucleotide-only linear model. Cbf1 was found to exhibit significant interactions between several pairs of flanking nucleotides^17^.

We instead used DFIM to directly query the Cbf1 neural network model and estimate pairwise nucleotide-resolution *FIS* between all pairs of nucleotides at all positions for all sequences in the library (Fig 8 B). We computed aggregate statistics (mean) of the absolute nucleotide-resolution FIS for all pairs of nucleotide features across the 5,000 sequences with strongest binding affinity (lowest measured ΔΔ G). We obtain four (40 x 40) aggregate DFIMs where each map corresponds to one of the 4 bases {*A, C, G, T*} as the observed source nucleotide. The rows in each 40 x 40 aggregate DFIM correspond to 4 mutant bases x 10 source positions, while the columns correspond to 4 target bases x 10 target positions. To ease interpretation, we compute a marginalized 40 x 40 DFIM that records the maximal average score over all mutant bases for each source base, source position, target base and target position (Fig 8 C), marginalized over the 3 potential mutations for a given source base. We observe that the marginalized aggregate DFIM for the high binding affinity sequences exhibit several strong interactions between flanking nucleotides (Fig 8 C).

**Figure 7:**
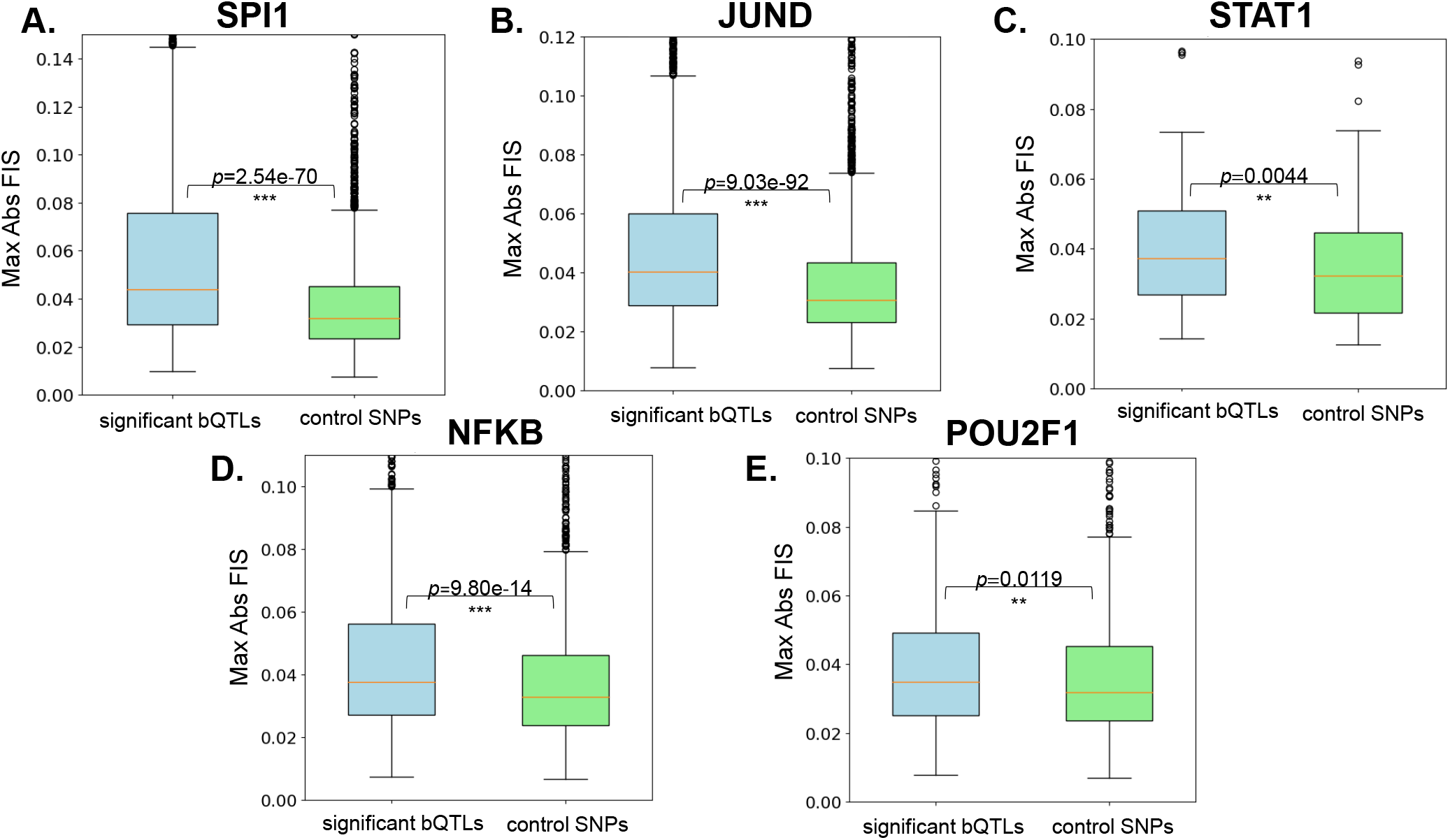
Distribution of maximal absolute feature interaction scores (*FIS*) for TF bQTLs and control SNPs for SPI1 (**A.**), JUND (**B.**), STAT1 (**C.**), NFKB (**D.**) and POU2F1 (**E.**). bQTLs of all TFs exhibit significantly stronger max *FIS* compared to control SNPs.

The map corroborates several of the strongest interactions identified by Le and Shimko using the distillation approach such as the strong interaction between a T at the −1 position and an A at the +1 position. Our maps also identify novel interactions such as a strong interaction between T at −1 and T at +2. In contrast, the aggregate DFIMs across 5,000 sequences with weakest binding affinity (highest measured ΔΔ G) exhibit uniformly weak interaction scores.

## 4. Discussion

We present an efficient method called Deep Feature Interaction Maps (DFIM) to identify epistatic interactions between all pairs of nucleotides or motif features in any DNA sequence input to a deep learning model for regulatory genomics. Our method accurately recovers ground truth interacting motifs in simulated regulatory DNA sequences. When applied to deep learning models of *in vivo* TF binding, we recover known proximal interactions between motifs of interacting co-factors while also discovering long-range interactions between motifs as far as 75 bp apart. We interpret deep learning models trained on *in vitro* TF binding to discover extensive interactions between pairs of nucleotides in sequences flanking core TF binding motifs. Finally, we interpret deep learning models of *in vivo* chromatin accessibility to generate nucleotide-resolution interaction maps for non-coding regulatory sequences surrounding SNVs (bQTLs) that affect binding of transcription factors. Our maps link binding QTLs to nearby sequence features including high and low affinity matches to the canonical binding site of the TF whose binding is disrupted. We also find bQTLs interacting with motifs of multiple co-binding TFs. These epistatic interactions seem to capture both cooperation and competition. While our primary focus in this manuscript is on interpreting feature interactions in DNA sequence inputs, DFIM can easily be generalized to other data modalities.

Partial dependence plots are commonly used to understand the sensitivity of a prediction to a one or more features18. DFIM serves as complementary approach to understand the predictive higher-order, non-linear interactions between features. DFIM is most efficient to estimate all pairwise interactions between pre-determined features such as known binding sites or SNVs or a sparse set of de-novo discovered predictive features with significant importance scores. However, DFIM also scales well to estimate interactions between all nucleotides in large sets of sequences because it leverages efficient backpropagation-based feature attribution methods. While DFIM is generally compatible with any efficient feature attribution method, we have not evaluated our approach on all such methods. However, we have found overall strong replication of DFIM results and associated conclusions by using two separate importance scores, namely DeepLIFT and gradient saliency maps. This suggests that DFIM could generalize to other importance scoring approaches.

**Figure 8:**
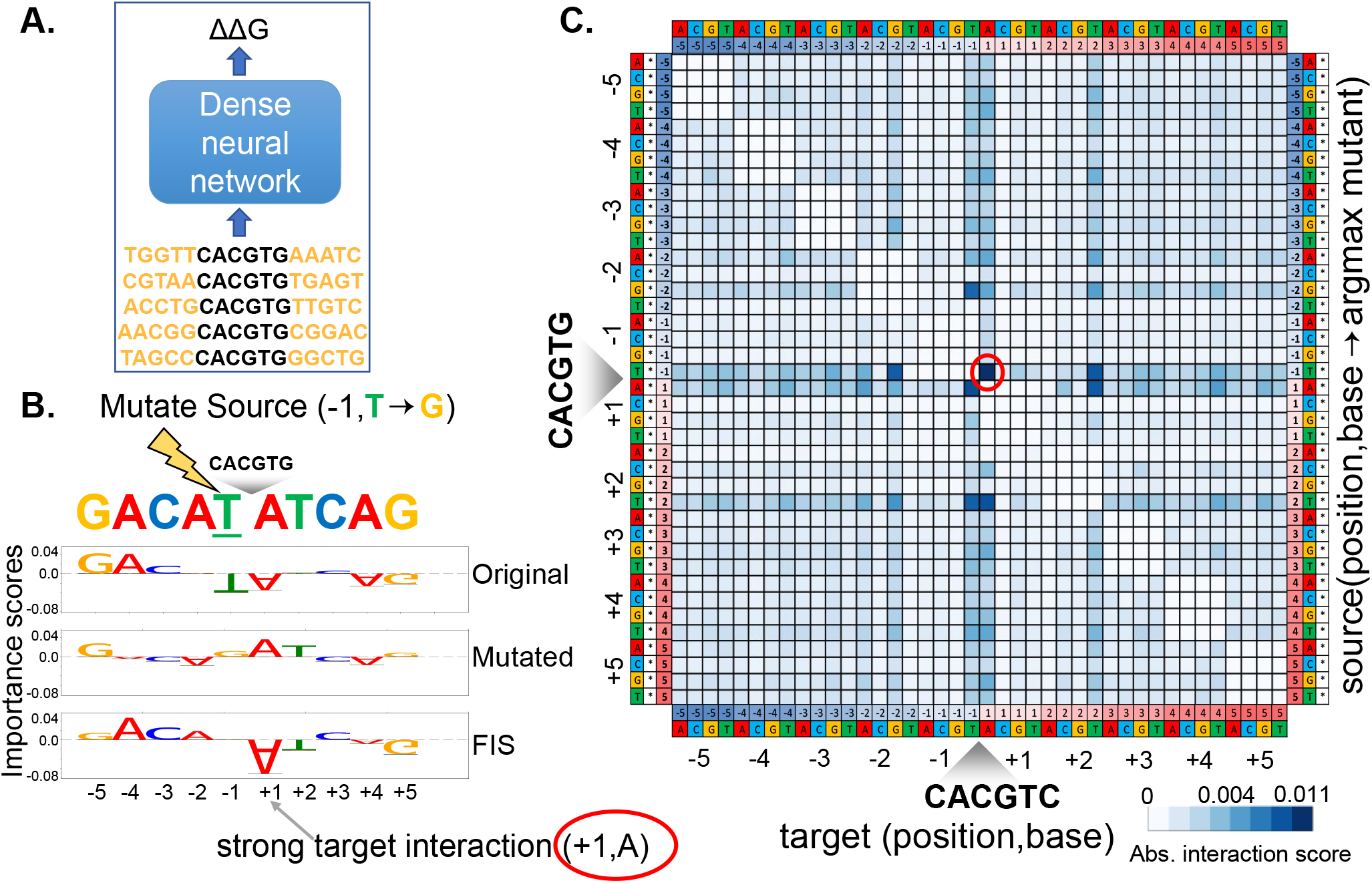
**A.** A feed forward neural network to fit binding affinities of a library of 10 bp sequences flanking the core Cbf1 motif. **B.** Source nucleotide A at source position −1 in an example sequence is mutated to a G. Row 1 and 2 show the importance scores of all nucleotides in the original and mutated sequence respectively. Row 3 shows the feature interaction scores (*FIS*) of all target nucleotides with respect to the source feature feature. We observe a strong interaction between source (−1, T) and target (+1, A). **C.** Marginalized aggregate deep feature interaction map (DFIM) for Cbf1 averaged across the top 5K highest binding affinity sequences. The rows correspond to (source position, source base, argmax mutant base). The columns correspond to (target position, target base). We observe a consistent strong interaction between source feature (−1,T) and target feature (+1,A).

There are some caveats to interpreting feature interactions derived from DFIM. Feature importance scores from any feature attribution method are only meaningful for examples that are predicted correctly. Since feature interaction scores from DFIM are based on feature importance scores, the validity of DFIM is also restricted to examples that are correctly predicted by high performance models. Further, vulnerabilities of the feature attribution method used in DFIM transfer over to the interaction scores. Hence, we recommend using multiple feature attribution methods to obtain robust estimates of interactions. Changes in model architecture can also change the interactions encoded by the model and thus the interactions learned with DFIM. Despite these mentioned caveats, the case studies we present here showcase the utility of DFIM to provide a nuanced view into the combinatorial code of regulatory DNA sequences through the lens of predictive neural network models.

## Acknowledgements

We would like to thank Johnny Israeli and Nathan Boley for their help training a TAL1, GATA1, GATA2 transcription factor binding model. We would also like to thank Avanti Shrikumar for discussions during early development of the methodology.

## Funding

PG was supported by a BioX Stanford Interdisciplinary Graduate Fellowship (SIGF). TS was supported by a National Science Foundation Graduate Research Fellowship. AK was supported by National Institute of Health grants 1DP2GM123485, 1U01HG009431 and 1R01HG00967401.

## References

1. Alipanahi B, Delong A, Weirauch MT, Frey BJ. 2015. Predicting the sequence specificities of DNA- and RNA-binding proteins by deep learning. Nat Biotechnol 33(8):831–838.

2. Kelley DR, Snoek J, Rinn JL. 2016. Basset: learning the regulatory code of the accessible genome with deep convolutional neural networks. Genome Res 26(7):990–999.

3. Zhou J, Troyanskaya OG. 2015. Predicting effects of noncoding variants with deep learning-based sequence model. Nat Methods 12(10):931–934.

4. Shrikumar A, Greenside P, Kundaje A. 2017. Learning Important Features Through Propagating Activation Differences. arXiv:1704.02685

5. Lundberg S, Lee SI. 2017. A Unified Approach to Interpreting Model Predictions. arXiv:1705.07874

6. Finnegan A, Song JS. 2017. Maximum entropy methods for extracting the learned features of deep neural networks. PLoS Comput Biol 13(10):e1005836.

7. Simonyan K, Vedaldi A, Zisserman A. 2013. Deep Inside Convolutional Networks: Visualising Image Classification Models and Saliency Maps. arXiv :1312.6034

8. Sundararajan M, Taly A, Yan Q. 2016. Gradients of Counterfactuals. arXiv:1611.02639

9. Lundberg SM, Erion GG, Lee SI. 2018. Consistent Individualized Feature Attribution for Tree Ensembles arXiv :1802.03888

10. Kheradpour P, Kellis M. 2014. Systematic discovery and characterization of regulatory motifs in ENCODE TF binding experiments. Nucleic Acids Res 42(5):2976–2987.

11. Gerstein MB, Kundaje A, Hariharan M, Landt SG, Yan KK, Cheng C, Mu XJ, Khurana E, Rozowsky J, Alexander R, Min R, Alves P, Abyzov A, Addleman N, Bhardwaj N, Boyle AP, Cayting P, Charos A, Chen DZ, Cheng Y, Clarke D, Eastman C, Euskirchen G, Frietze S, Fu Y, Gertz J, Grubert F, Harmanci A, Jain P, Kasowski M, Lacroute P, Leng J, Lian J, Monahan H, O’Geen H, Ouyang Z, Partridge EC, Patacsil D, Pauli F, Raha D, Ramirez L, Reddy TE, Reed B, Shi M, Slifer T, Wang J, Wu L, Yang X, Yip KY, Zilberman-Schapira G, Batzoglou S, Sidow A, Farnham PJ, Myers RM, Weissman SM, Snyder M. 2012. Architecture of the human regulatory network derived from ENCODE data. Nature 489(7414):91–100.

12. ENCODE Project Consortium. 2012. An integrated encyclopedia of DNA elements in the human genome. Nature 489(7414):57–74.

13. Kassouf MT, Hughes JR, Taylor S, McGowan SJ, Soneji S, Green AL, Vyas P, Porcher C. 2010. Genome-wide identification of TAL1’s functional targets: insights into its mechanisms of action in primary erythroid cells. Genome Res 20(8):1064–1083.

14. Behera V, Evans P, Face CJ, Hamagami N, Sankaranarayanan L, Keller CA, Giardine B, Tan K, Hardison RC, Shi J, Blobel GA. 2018. Exploiting genetic variation to uncover rules of transcription factor binding and chromatin accessibility. Nat Commun 9(1):782.

15. Tehranchi AK, Myrthil M, Martin T, Hie BL, Golan D, Fraser HB. 2016. Pooled ChIP-Seq Links Variation in Transcription Factor Binding to Complex Disease Risk. Cell 165(3):730–741.

16. Corces MR, Trevino AE, Hamilton EG, Greenside PG, Sinnott-Armstrong NA, Vesuna S, Satpathy AT, Rubin AJ, Montine KS, Wu B, Kathiria A, Cho SW, Mumbach MR, Carter AC, Kasowski M, Orloff LA, Risca VI, Kundaje A, Khavari PA, Montine TJ, Greenleaf WJ, Chang HY. 2017. An improved ATAC-seq protocol reduces background and enables interrogation of frozen tissues. Nat Methods 14(10):959–962.

17. Le DD, Shimko TC, Aditham AK, Keys AM, Longwell SA, Orenstein Y, Fordyce PM. 2018. Comprehensive, high-resolution binding energy landscapes reveal context dependencies of transcription factor binding. Proc Natl Acad Sci U S A 115(16):E3702–E3711.

18. Friedman JH. 2001. Greedy function approximation: A gradient boosting machine. The Annals of Statistics 29(5):1189–1232.

